# *In silico* agent-based modeling approach to characterize multiple *in vitro* tuberculosis infection models

**DOI:** 10.1101/2023.03.13.532338

**Authors:** Alexa Petrucciani, Alexis Hoerter, Leigh Kotze, Nelita Du Plessis, Elsje Pienaar

## Abstract

*In vitro* models of *Mycobacterium tuberculosis (Mtb)* infection are a valuable tool to examine host-pathogen interactions and screen drugs. With the development of more complex *in vitro* models, there is a need for tools to help analyze and integrate data from these models. We introduce an agent-based model (ABM) representation of the interactions between immune cells and bacteria in an *in vitro* setting. This *in silico* model was used to independently simulate both traditional and spheroid cell culture models by changing the movement rules and initial spatial layout of the cells. These two setups were calibrated to published experimental data in a paired manner, by using the same parameters in both simulations. Within the calibrated set, heterogeneous outputs are seen for outputs of interest including bacterial count and T cell infiltration into the macrophage core of the spheroid. The simulations are also able to predict many outputs with high time resolution, including spatial structure. The structure of a single spheroid can be followed across the time course of the simulation, allowing the relationship between cell localization and immune activation to be explored. Uncertainty analyses are performed for both model setups using latin hypercube sampling and partial rank correlation coefficients to allow for easier comparison, which can provide insight into ideal use cases for the independent setups. Future model iterations can be guided by the limitations of the current model, specifically which parts of the output space were harder to reach. This ABM can be used to represent more *in vitro Mtb* infection models due to its flexible structure, providing a powerful analysis tool that can be used in tandem with experiments.

**Author Summary:** Tuberculosis is an infectious disease that causes over 1.4 million deaths every year. During infection, immune cells surround the bacteria forming structures called granulomas in the lungs. New laboratory models generate spheroids that aim to recreate these structures to help understand infection and find new ways to treat tuberculosis. Computational modeling is used to compare these newer spheroid models to traditional models, which don’t recreate the structure of the cell clusters. After calibration to data from laboratory experiments to ensure that the computational model can represent both systems, the structures were characterized over time. The traditional and spheroid model were also compared by looking at how model inputs impact outputs, allowing users to figure out when one model should be used over the other. This computational tool can be used to help integrate data from different laboratory models, generate hypothesis to be tested in laboratory models, and predict pathways to be targeted by drugs.

## 1. Introduction

Tuberculosis (TB) continues to be a global public health crisis, responsible for 1.4 million deaths in 2021 alone.(1) TB is caused by the bacteria *Mycobacterium tuberculosis* (*Mtb*). Generally, *Mtb* is introduced to its host upon inhalation of contaminated respiratory droplets, allowing direct entry into the lungs. Bacteria are deposited in the well-ventilated lower lobes of the lung, where alveolar macrophages phagocytose them.(2) *Mtb* is subsequently able to survive and replicate within the endosomes of these macrophages.(3) As the infection progresses, infected macrophages release chemokines and cytokines which recruit other immune cells (e.g. monocytes, T cells, B cells, NK cells, dendritic cells, and neutrophils) to form a granuloma. A granuloma is generally comprised of a core of infected macrophages, surrounded by monocytes, epithelioid macrophages, foamy macrophages, neutrophils, multinucleated giant cells, and finally a lymphocytic cuff with an outer fibrous capsule.(4) The timing and spatial organization of key host-pathogen interactions within these granuloma structures, and how these interactions contribute to bacterial survival or elimination, remains incompletely understood. This is in part due to the complexity of the granuloma structure itself, which makes it difficult to understand, measure, and/or predict host-pathogen interactions and their impact on infection progression.

Many systems have been used to explore granulomas in TB; each having its own benefits and limitations. While much has been revealed about the structure of granulomas from work in humans, clinical studies are invasive or indirect and are often lacking in time points required to evaluate granuloma dynamics. Additionally, TB granulomas in humans can only really be studied at later stages when the infection has been established and diagnosed.(5) Animal studies such as non-human primate (NHP), rabbit, and mouse models are very useful and allow more control and direct observation of infection and granuloma formation than in humans. Mouse models benefit from wide availability of commercial immunological reagents, genetic tools, and transgenic and knock-out strains, but most mouse strains struggle to recreate the structure of granulomas seen in humans.(6, 7) Rabbit and guinea pigs are able to form necrotic and non-necrotic mature granulomas. (6, 7) These models have been limited in the past by availability of immunological reagents, but recently more commercially available immunological reagents like antibodies against rabbit analytes have been developed.(6–9) NHP models most closely recreate human pathology, with heterogenous clinical outcomes and granuloma structures.(10, 11) But NHP models are expensive, time-intensive, and limited by the availability of animal facilities.(6, 7) It is difficult to do certain genetic manipulations, collect data at many time points, and control the exact cellular and environmental makeup of the system in these *in vivo* models. Complementary to these *in vivo* models, there has been recent work developing more complex *in vitro* cellular cultures to both dissect biological mechanisms and test new therapies (reviewed in Elkington et al.(12)). *In vitro* models can be particularly helpful because the system is tractable, and all cellular components of the system can be controlled. *In vitro* models are also cheaper and higher throughput than the equivalent *in vivo* models. *In vitro* systems can be mechanistically perturbed and dynamically sampled in ways that are extremely difficult in *in vivo* models.

Elkington et al. suggest certain criteria for an ideal *in vitro* model including the use of human cells and virulent *Mtb*; allowing incorporation of fibroblast, epithelial cells, and physiological extracellular matrix; being modular to allow many different biological questions to be answered; and, ideally, being 3-dimensional (3D).(12) However, increasing complexity isn’t necessary in all cases and can make models lower-throughput and more expensive. Ideally, *in vitro* models could be tailored to the biological question at hand, but still be able to be compared across platforms. *In vitro* models could then be optimized to include only the necessary components, allowing maintenance of inexpensive, high-throughput models. Results from many disparate systems could still be synthesized to form robust conclusions.

We recently developed an *in vitro* biomimetic 3D spheroid granuloma model.(13) Briefly, patient-derived alveolar macrophages are infected with BCG, and magnetic nanospheres used to levitate the cells. Autologous adaptive immune cells isolated from peripheral blood mononuclear cells (PBMCs) were added at 48 hours into the 6 day culture. When comparing this granuloma model to a corresponding traditional monolayer culture, we found the spheroid model was better able to control bacteria. Differences in bacterial count between these models can be quantified and are due to the different model setups, but how the spatial aspects impact immune response is unclear. These two systems provide a good test case to evaluate the possibility of translating between different *in vitro* systems, and identify the key mechanisms at work in the different systems.

This data not only motivates a need to understand the mechanistic differences between these two models, but also highlights a need to more broadly look at the complexity and spatiality of *in vitro* models. As we move towards more complex *in vitro* models, organoids, complex cell mixtures etc., it is important that we 1) understand and quantify the impact of the structural organization of the cultures, and 2) develop tools that are able to analyze these more complex systems, and 3) develop tools that can enable us to compare and translate between systems. Computational models are well-suited to address all of these tasks.

Computational models are inexpensive compared to *in vitro* or *in vivo* models, quick to run, highly manipulatable, able to integrate data from many sources, and can easily be adapted to reflect new data.(12, 14) Beyond this, computational models can be used to perform perturbations (e.g. virtual knockouts) that would be extremely difficult in a wet lab setting. Computational models work especially well in combination with *in vitro* work, where hypotheses can be generated computationally and tested experimentally in an iterative fashion.(12) Mechanistic models specifically use individual interactions between cells and molecules to predict emergent tissue-level outcomes (e.g. granuloma dynamics). Because individual cellular and molecular interactions are based on current biological understanding, we can use the emergent behavior of our simulations to test hypotheses about the driving mechanisms for tissue-level outcomes. Beyond hypothesis testing, mechanistic models also act to integrate existing knowledge into a single framework to help understand their collective impact. One type of mechanistic model, agent-based models (ABMs), are stochastic spatiotemporal models that are particularly suited to look at emergent spatial behavior. Stochasticity is ideal because it captures some of the heterogenous host response to TB.(15, 16) Spatiality is required as we aim to represent and contrast both traditional and 3D models.

Mechanistic modeling has been applied to TB since 1962, and ABMs in particular have been used in the context of TB since 2004.(17–19) ABMs of granuloma formation in the non-human primate (NHP) lung have been iterated many times to look at the impacts of TNF-α(20–22), *Mtb* metabolism(23), macrophage (MΦ) polarization(24), and more(25–30). In this work, we apply these established agent-based approaches to *in vitro* systems. This means that all components included in the experimental system can be accounted for, the experimental system can be more easily observed and perturbed, and we can use one simulation framework with different initializations to represent, and translate between, many *in vitro* models. In this work we use one computational agent-based modeling framework to recreate the results from both 3D spheroid and the corresponding traditional culture in vitro models(13). Our computational model generates high time-resolution data for cellular outputs, along with spatial data. This spatial data is processed in multiple ways, allowing us to dissect the evolution of a single granuloma and explore the heterogeneity of the host response within different spatial organizations. Finally, we use uncertainty analysis to look at the similarities and differences between the spheroid and traditional setups.

## 2. Methods

### 2.1. Experimental Methods

The data we use for calibration is derived from a biomimetic 3D spheroid model of a granuloma and the corresponding traditional culture. Briefly, HIV negative patients with high suspicion of TB were recruited. Bronchoscopies were performed by qualified clinicians and nursing staff according to international guidelines (31) to obtain bronchoalveolar lavage fluid samples. Immediately after bronchoscopy, peripheral blood was collected by venipuncture into two 9mL sodium heparinized (NaHep) vacutainers. Alveolar macrophages were isolated from bronchoalveolar fluid, and PBMCs were isolated from peripheral blood using the Ficoll-Paque isolation method described previously (13). Alveolar macrophages were cultured at a density of 4×10^5^ cells per well in a 24-well low-adherence culture plate and infected with *Mycobacterium bovis* Bacille Calmette-Guerin (BCG) at a multiplicity of infection (MOI) of 1 for 4 hours. Afterwards, extracellular bacteria were removed by supplementing media with an antimycotic antibiotic (penicillin/streptomycin/amphotericin B) for 1 hour, followed by successive washes. The 3D spheroids were made by treating alveolar macrophages with biocompatible NanoShuttle (n3D Biosciences Inc., Greiner Bio-One) and levitating them using the magnetic levitating drive. After 48 hours, 6×10^5^ autologous CD3+ T cells are added per well. The traditional culture is made using the same cells and the same ratios, but without NanoShuttle treatment and subsequent magnetic levitation.

Granuloma structures were mechanically disrupted by gentle pipetting after 6 days of culture. Cell count and cell viability were determined using the trypan blue exclusion method after adherent cells were removed. CFU counts were determined by lysing mechanically disrupted cells and plating serial dilutions on Middlebrook 7H11 agar plates (BD Biosciences). After 21 days of growth, the colonies were manually counted. 3D spheroids were also fixed, embedded in tissue-freezing medium OCT (Tissue-Tek; USA), and cryosectioned. A section from the middle of the structure was stained with antibodies for CD3+ and CD206+ cells and imaged using a Carl Zeiss LSM 880 Airyscan with Fast Airyscan Module confocal microscope (Plan-Apochromat x63/1.40 oil DIC UV-VIS-IR M27 lens objective). The image of the traditional cell culture was acquired with light microscopy at 40x magnification. For full methods please reference Kotze et al. 2021.

### 2.2. Model Structure

Our model simulates the interactions between macrophages, CD4+ T cells, CD8+ T cells, bacteria, and two simplified cytokines within an *in vitro* environment. The simulation is constructed as a hybrid multiscale model with a cellular level agent-based model hybridized to a partial differential equation model of diffusion for the two cytokines (TNFα-like and IFNγ-like). These will be referred to as TNFα and IFNγ moving forward. The environment is composed of grid cubes that each represent a 20μm × 20μm × 20μm volume, which is the approximate size of our largest agent type, the macrophage.(32) The environment has two overlying grids, one single occupancy grid for immune cells and one multioccupancy for the smaller bacteria. The simulation has 4 types of agents: macrophages, CD4+ T cells, CD8+ T cells, and bacteria. Macrophages can be subdivided into uninfected and infected classes. Agent behaviors are performed with a time step of 6 minutes, the approximate time for a monocyte to move 20 μm, or one grid cube. (33–36) The simulation is run for a total of 6 days, to reflect the duration of the *in vitro* experiments. An overview of agent behaviors is shown in Figure 1 and further detail is given below. These methods are in part drawn from work modeling NHP granulomas *in silico*, specifically GranSim and subsequent developments.(18, 19)

**Figure 1:**
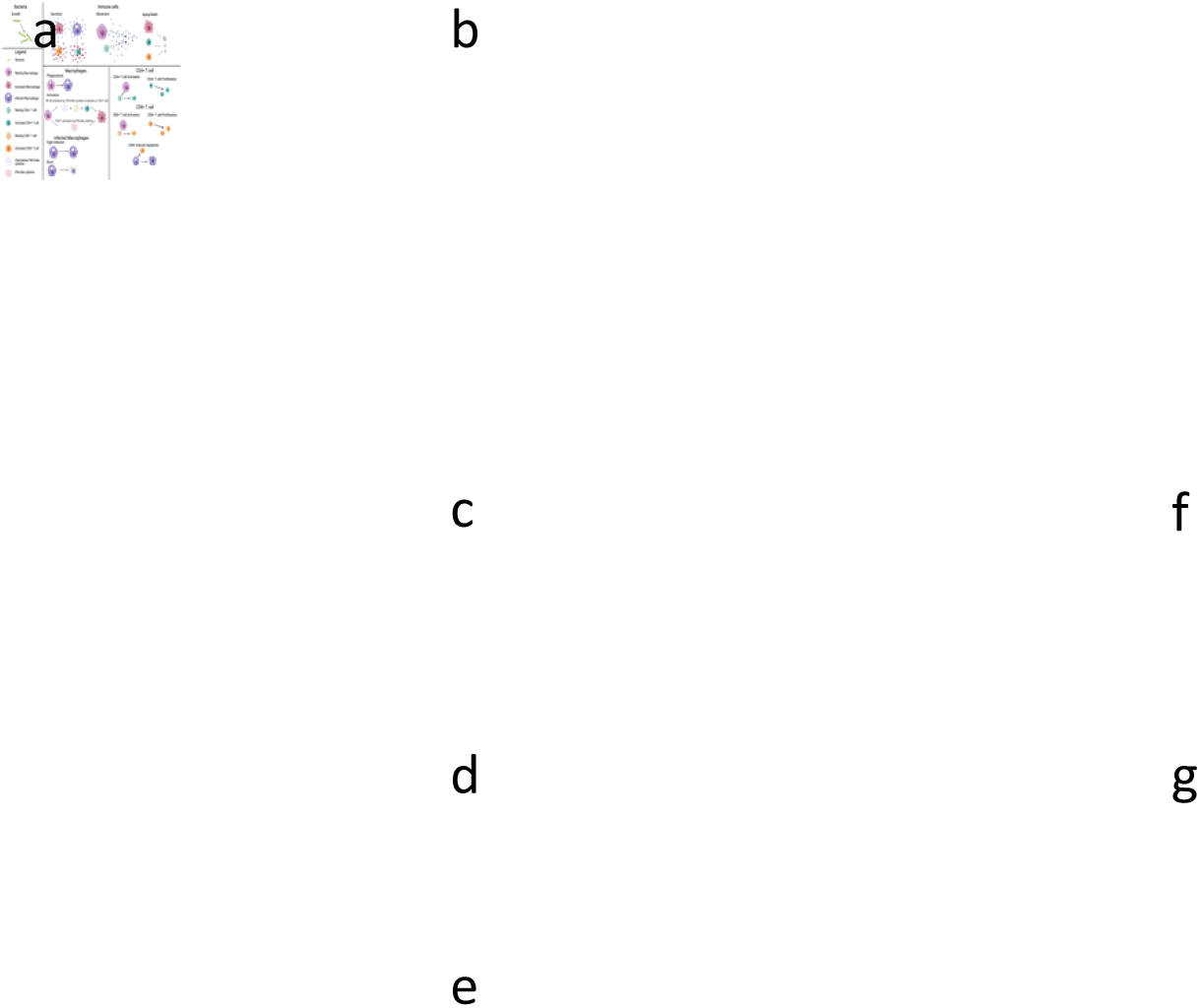
An overview of rules for the simulated agents. a) Bacteria grow and divide. b) Immune cells secrete cytokines dependent on activation or infectious state, move probabilistically up a TNF-α gradient, age, and die. c) Macrophages (MΦ) can phagocytose bacteria becoming infected. d) MΦ activation is represented by a two-step process. NF-κB can be activated by TNF-α, bacteria, or direct contact with an activated CD4+ T cell. STAT1 is activated by IFN-γ secreted by activated T cells. e) Infected ΜΦ either fight infection killing internal bacteria and returning to uninfected state, or when a certain threshold of bacteria is reached will burst releasing internal bacteria into the environment. f) TB-Specific CD4+ T cells activate by interacting with a ΜΦ that has interacted with a bacterium. After activation, CD4+ T cells can proliferate. g) TB-Specific CD8+ T cells activate by interacting with a ΜΦ that has interacted with a bacterium and is STAT1 activated. After activation, CD8+ T cells can proliferate and kill infected macrophages along with the internal bacteria. Created with BioRender.com.

#### 2.2.1. Diffusing molecules

There are 2 diffusing molecules included representing the simplified TNF-α and IFN-γ. These are contributed to by the secreting agents, and diffuse in the simulation space. Diffusion is performed similarly to that in Weathered et al. using a 3D alternating-direction explicit numerical method.(37) As this method is unconditionally numerically stable, a larger *dt* than is predicted by the conditional stability criterion can be used while maintaining accuracy.(38) After finding *dt* suggested by the conditional stability criterion and the diffusion parameters a multiplier of 4 was incorporated into the alternating-direction explicit method to reduce simulation time, while maintaining accuracy, as recommended by Cilfone et al.(38) The PDE is run with a smaller time step than the ABM, ranging from 2 to 14 diffusion iterations per agent time step depending on the diffusion parameters. IFN-γ and TNF-α are diffused separately with separate diffusion coefficients and decay rates. The rate of diffusion is slowed within granulomas by *granulomaFractionOfDiffusion*.

#### 2.2.2. Agents

##### Immune cells

Macrophages, infected macrophages, CD4+ and CD8+ T cells are all classified as types of immune cells. This parent class of agents share common behaviors, including movement and aging. Movement is determined by gravity limited or 3D rules. Cells moving in 3D are able to move in any direction. With gravity limited rules, cells will fall in the z dimension if no immune cell is below them and can only move up in the z direction if on top of another immune cell. Given these movement rules, the cells will chemotax probabilistically toward the highest concentration of TNF-α when the summed TNF-α in the Moore neighborhood is above *TNFthresholdForImmuneCellMovement*. This chemotaxis algorithm is based off of that in Weathered et al..(37) Immune cells also age according to individualized lifespans. A resting lifespan and activated lifespan are selected for each cell from a *populationLifespan* * (1+/- *lifeSpanVariance*). These lifespans are then converted to aging rates, which change according to the activation status of the cell. The resting aging rate is 1 hour aged per hour, while the activated aging rate is calculated as resting lifespan divided by activated lifespan. At initialization a cell will be given a random starting age from zero to the resting lifespan. Then a cell’s current age gets incremented by the aging rate each time step. When a cell reaches its maximum age, it will die and be removed from the simulation.

##### Macrophages

Beyond the immune cell rules described above, macrophages will attempt to phagocytose and activate every time step. Each macrophage attempts to phagocytose by picking a bacterium in its Moore neighborhood at random. If this bacterium is extracellular, it will be phagocytosed with a phagocytosis probability dependent on activation state (*basePhagocytosisProbability*, *activePhagocytosisProbability*). Successful phagocytosis turns a macrophage into an infected macrophage. Macrophages that have phagocytosed bacteria also get classified as having interacted with bacteria, meaning antigenic peptides can be displayed on the cell surface. Each macrophage also checks for activation. Activation is represented by a simplified two step signaling process, requiring STAT1 and NF-κB activation.(39) Each of these two pathways can be activated, if they are not already activated. STAT1 is activated if local IFN-γ is greater than *IFNthresholdForStat1Activation*. NF-κB can be activated in 3 ways: TNF-α greater than *TNFthresholdForNFkBActivation*, nearby extracellular bacteria greater than *bacThresholdForNFkBActivation*, or direct interaction with an activated CD4+ T cell. These represent TNF-α interaction with TNFR, activation of TLR, and CD40-CD40L interactions, respectively.(40) All three of these NF-κB activation methods will be checked in a random order. NF-κB and STAT1 activations last for set durations after the signal was initially received (*nfkbSpan* and *stat1Span*). These durations have variances, *nfkbVariance* and *stat1Variance*, to introduce heterogeneity into the population. After the macrophage-specific length of activated time, the pathway will deactivate and be checked again immediately, to allow longer activation if the activation signals persist. If both pathways are activated at the same time, then the macrophage becomes fully activated. Activation changes a macrophage’s movement probability, phagocytosis probability, and aging rate. Activated macrophages also secrete TNF-α at a rate of *ActivatedMacrophageTNFSecretion* molecules per second.

##### Infected Macrophages

Infected macrophages can fight the infection at each time step. An internal bacterium is selected randomly and will be killed with a probability that is dependent on the macrophage’s activation state (*baseKillingProbability*, *activeKillingProbability*). If all the bacteria within an infected macrophage are killed, then the infected macrophage reverts to a healthy macrophage. Infected macrophages can be activated through the same pathways as healthy macrophages. When fully activated, the phagocytosis and killing probabilities change to values for activated macrophages. Infected macrophages secrete TNF-α when activated, but also constitutively secrete TNF-α at a baseline level of *InfectedMacrophageTNFSecretion* molecules per second when not activated. Infected macrophages don’t move but can continue to phagocytose bacteria if the number of internalized bacteria is below *phagocytosisThreshold*. This occurs similarly to the initial phagocytosis, with a random bacterium selected from the infected macrophage’s Moore neighborhood that will be taken up with some probability if it is extracellular. Once the number of internal bacteria is above *cellularDysfunctionThreshold* the macrophage is considered chronically infected.(19) Chronically infected macrophages can no longer be fully activated or kill internal bacteria. If the number of bacteria within an infected macrophage reaches a bursting threshold the macrophage will burst and release the internal bacteria into the environment. This threshold has been experimentally determined to be 20-40 internal bacteria in *in vitro* systems.(41) A burst limit was randomly selected for each infected macrophage from a uniform distribution from 20 to 40 internal bacterial. When a macrophage dies of old age the bacteria are similarly released into the environment.

##### CD4+ T cells

CD4+ cells can be TB specific or non-TB specific. TB specific CD4+ T cells can also become activated. Activation of TB specific CD4+ T cells occurs with a probability of *CD4ActivationProbability* if a random macrophage in its Moore neighborhood has interacted with bacteria. This is equivalent to antigen presentation on MHC II. (40) Activation increases movement probability and aging rate. Activated CD4+ T cells secrete both TNF-α at *ActivatedCD4TNFSecretion* molecules per second and IFN-γ at *ActivatedCD4IFNSecretion* molecules per second.(42) Active CD4+ T cells can also divide with a doubling time of *cd4PopulationDoublingTime* until the maximum number of generations (*maximumCD4Generations*) is reached. Individual variance is introduced to doubling time.

Deactivation occurs with a given probability *CD4DeactivationProbability* per time step.

##### CD8+ T cells

Just like CD4+ T cells, CD8+ T cells can be subdivided into TB specific and non-TB specific. TB specific CD8+ T cells can be activated. If a randomly selected macrophage within the T cell’s Moore neighborhood is STAT1 activated and has interacted with bacteria, then the T cell will probabilistically activate (*CD8ActivationProbability*). STAT1 activation is a proxy for interaction between CD4+ T cell and macrophage which increases expression of molecules on the surface of the APC(B7 and 4-1BBL) that provide co-stimulation to naïve CD8+ T cells.(40, 43) If activated, a CD8+ T cell will secrete both TNF-α (*ActivatedCD8TNFSecretion*) and IFN-γ (*ActivatedCD8IFNSecretion*). Activation also increases movement probability and aging rate. Activated CD8+ T cells will also divide with a doubling time of *cd8Population_DoublingTime* until the maximum generation (*maximumCD8Generations*) is reached. Activated CD8+ T cells have the ability to kill infected macrophages (equivalent to cells presenting peptides in MHC I). A random infected macrophage is selected for the Moore neighborhood, and the infected macrophage and all internal bacteria are killed with a probability *CD8KillProbability*. CD8+ T cells deactivate probabilistically (*CD8DeactivationProbability*).

##### Bacteria

Bacteria grow and divide. Bacteria have biomass that gets added to every tick. The rate of growth depends on whether they are intracellular or extracellular. Growth rate is calculated from doubling time (*mtbInternalDoublingTime*, *mtbExternalDoublingTime*), and includes some individual variance from the population mean. If the biomass threshold of 2 is reached, then the bacteria divide into two with the biomass distributed among them unevenly(44). Simulated bacteria represent BCG, as BCG was used in the *in vitro* models. Behaviors/parameters draw from both BCG and TB literature.

#### 2.2.3. Initial Conditions

The differences between the spheroid and traditional simulations include the movement rules and the initial spatial distribution of cells. Our initial conditions reflect those used in the *in vitro* system.(13)

##### Spheroid

In the experimental protocols, 400,000 macrophages are infected with MOI 1 and then levitated.(13) At day 2, 600,000 CD3+ cells are added in a dropwise manner directly to the spheroid. Due to computational limitations associated with the 3D simulation of a full-sized spheroid, we simulate a spheroid of 1/10^th^ the size. We generate a sphere of 40,000 mixed healthy and infected macrophages. Given the experimental MOI of 1, we use a Poisson distribution to estimate percentage of cells with various number of phagocytosed bacteria(45). The fraction of macrophages that have phagocytosed *n* bacteria is given by 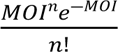 Macrophages with zero to six internalized bacteria are initialized, giving 39,997 initial bacteria. This sphere is centered on an 80×80×80 grid representing 1.6 mm × 1.6 mmx 1.6 mm volume. The radius of the initialized sphere is calculated as 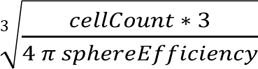, with the initial density of the cells determined by *sphereEfficiency*. At day 2, 60,000 CD4+ and CD8+ T cells are added in a cuff around the macrophages. Proportions of CD4+ T cells (*fractionCD4*), CD8+ T cells (*CD8Fraction*), and TB specific T cells (*fractionTBSpecific*, *tbSpecificCD8Fraction*) are estimated from literature. (46–49) Subsets of the immune cells are allowed to be preactivated (*activatedMacrophageProportion*, *activatedTBSpecificCD4Fraction*, *activatedTBSpecificCD8Fraction*) as the alveolar macrophages and PBMCs were taken from patients with active TB. Activated TB specific T cells are given a random starting generation and starting point in the division cycle as the process of proliferation could have already started.

##### Traditional culture

The experimental conditions are the same as the spheroid without the inclusion of the magnetic levitation beads. As with the spheroid, a simulation 1/10^th^ the size of the experiment. This is simulated by adding 40,000 infected and uninfected macrophages distributed evenly through the environment. After these macrophages are added they fall to the bottom of the plate due to the gravity-limited movement discussed in section 2.2.2.1. Since the cells would all be at the bottom of the plate, the dimensions were adjusted to 216×216×11, or 4.32mmx4.32mmx0.22 mm. The ratio of cells to the surface area of the plate is kept constant between the experimental system and the simulation. Additionally, the volume of simulation, and therefore initial cellular density, is minimally different between the spheroid and traditional models. The percentage of cells with various number of phagocytosed bacteria is calculated in the same manner as the spheroid model. On day 2, 60,000 CD4+ and CD8+ T cells are distributed evenly throughout the environment before falling.

#### 2.2.4. Simulation

This model is built using Repast Simphony 2.8, an open source software used to build ABMs in Java.(50) Simulations were run on the Purdue Brown Cluster and on XSEDE resources.(51) Python and MATLAB were used for data analysis and visualization.

### 2.3. Calibration

Calibration is performed by doing an initial parameter sweep and then iterating around specific parameter sets. These iterations are used to find a variety of parameter sets that fit into the experimental data range while iterating into harder to reach parts of the output space.

Experimental data ranges used for calibration include:

- Spheroid bacterial fold change from 4 hpi to day 6
- Traditional bacterial fold change from 4 hpi to day 6
- Spheroid cell viability at day 6
- Traditional cell viability at day 6
- Spheroid cell count at day 6
- Traditional cell count at day 6

A total of 50 parameters are varied in the model (Table 1). Initial ranges are determined from relevant literature (*in silico*, *in vivo*, *in vitro*) or left broad. Latin hypercube sampling (LHS) was used to sample 1,000 parameter sets from initial ranges with a centered design (Table 1).

**Table 1:** Parameters that are varied during calibration. Initial ranges are either determined by literature, estimated through preliminary simulations (e), or broadened to the full mathematically possible range (f). The set of calibrated parameter sets can be found in the provided data.

| Parameter | Initial Range | Units | Refs |
| --- | --- | --- | --- |
| **Bacteria** |  |  |  |
| *mtbInternalDoublingTime* | 23,69 | Hours | (53) |
| *mtbExternalDoublingTime* | 23,69 | Hours | (53) |
| **Macrophages** |  |  |  |
| *activatedMacrophageProportion* | 0,0.1 | Per tick | e |
| *baseKillingProbability* | 0.0001,0.02 | Per tick | e |
| *activeKillingProbability* | 0.002,0.3 | Per tick | e |
| *basePhagocytosisProbability* | 0,1 | Per tick | f |
| *activePhagocytosisProbability* | 0,1 | Per tick | f |
| *phagocytosisThreshold* | 8,12 | Internal bacteria | (22) |
| *cellularDysfunctionThreshold* | 8,12 | Internal bacteria | (22) |
| *nfkbSpan* | 0.16,166 | Hours | (24) |
| *TNFthresholdForNFkBActivation* | 40,500 | Molecules | e |
| *bacThresholdForNFkBActivation* | 20,150 | External bacteria | (22) |
| *stat1Span* | 0.16,166 | Hours | (24) |
| *IFNthresholdForStat1Activation* | 40,500 | Molecules | e |
| *ActivatedMacrophageTNFSecretion* | 0,40 | Molecules/second | (42) |
| *InfectedMacrophageTNFSecretion* | 0,40 | Molecules/second | (42) |
| *macrophagePopulation_MaxLifespan* | 20,100 | Days | (22) |
| *macrophagePopulation_MaxActivatedLifespan* | 7,13 | Days | (22) |
| *baseMovementProbabilityMacro* | 0.5,1 | Per tick | (33–36) |
| *activatedMovementProbabilityMacro* | 0,0.5 | Per tick | e |
| **CD4+ T cells** |  |  |  |
| *fractionCD4* | 0.5,0.65 | CD4+ T cells/ CD3+ T cells | (46,47) |
| *fractionTBSpecific* | 0.0001,0.06 | TB specific CD4+ T cells/<br>Total CD4+ T cells | (48,49<br>) |
| *activatedTBSpecificCD4Fraction* | 0,0.1 | Initial activated TB<br>specific CD4 T cells/ Total<br>TB specific CD4 T cells | e |
| *CD4ActivationProbability* | 0,1 | Per tick | f |
| *CD4DeactivationProbability* | 0,1 | Per tick | f |
| *ActivatedCD4TNFSecretion* | 0,40 | Molecules/second | (42) |
| *ActivatedCD4IFNSecretion* | 0,40 | Molecules/second | (42) |
| *cd4PopulationDoublingTime* | 6,16 | Hours | (54,55<br>) |
| *maximumCD4Generations* | 3,10 | Generations | (54,56<br>,57) |
| *cd4Population_MaxLifespan*<br>*cd8Population_MaxLifespan* | 34,340 | Days | (58–<br>60) |
| *cd4Population_ActivatedLifespan*<br>*cd8Population_MaxActivatedLifespan* | 2.5,4 | Days | (22,54<br>) |
| *baseMovementProbabilityCD4*<br>*baseMovementProbabilityCD8* | 0,1 | Per tick | f |
| *activatedMovementProbabilityCD4*<br>*activatedMovementProbabilityCD8* | 0,1 | Per tick | f |
| **CD8+ T cells** |  |  |  |
| *CD8Fraction* | 0.3,0.35 | CD8+ T cells/ CD3+ T<br>cells | (46) |
| *tbSpecificCD8Fraction* | 0.0001,0.06 | TB specific CD8+ T cells/<br>Total CD8+ T cells | (48,49<br>) |
| *activatedTBSpecificCD8Fraction* | 0,0.1 | Initial activated TB<br>specific CD8 T cells/ Total<br>TB specific CD8 T cells | e |
| *CD8ActivationProbability* | 0,1 | Per tick | f |
| *CD8DeactivationProbability* | 0,1 | Per tick | f |
| *ActivatedCD8TNFSecretion* | 0,40 | Molecules/second | (42) |
| *ActivatedCD8IFNSecretion* | 0,40 | Molecules/second | (42) |
| *cd8PopulationDoublingTime* | 3,13 | Hours | (55) |
| *maximumCD8Generations* | 7,20 | Generations | (56,57<br>,61) |
| *CD8KillProbability* | 0.012,0.12 | Per tick | (22) |
| **Diffusion** |  |  |  |
| *TNFthresholdForImmuneCellMovement* | 1,500 | Molecules | e |
| *TNFDiffusionCoefficient* | 0.1,1 | $10^{-7} \text{ cm}^2/\text{s}$ | (24) |
| *TNFDegradationRatePerSecond* | 0.96,10 | 1/s | e |
| *IFNDiffusionCoefficient* | 0.1,1 | $10^{-7} \text{ cm}^2/\text{s}$ | (24) |
| *IFNDegradationRatePerSecond* | 0.96,10 | 1/s | e |
| *granulomaFractionOfDiffusion* | 0,1 | - | f |
| *sphereEfficiency* | 0.65,0.9 | - | e |

These parameter sets are run in both the traditional and spheroid simulation with 7 replicates as a broad initial sweep. Top runs are defined as those with the highest traditional CFU, as this part of the output space had few runs in the initial sweep. The top five runs that met the bacterial fold changes for traditional and spheroid are iterated. Iterations are performed by narrowing the parameter range to 20% of the initial range centered around the initial point (each of the top five runs). One hundred samples in this new range are generated using LHS and are run in triplicate. The number of replicates and runs are reduced due to computational costs.

Runs that passed all 6 criteria (bacterial fold changes, cell viability, and cell count at day 6 for traditional and spheroid cultures) are iterated until there was less than a 10% increase in traditional culture CFU. The iterating range is then narrowed to 10% of the initial range, and iterated until again there is a less than 10% increase in traditional culture CFU. The calibrated set is generated by selecting runs that fits all 6 criteria from all of the simulations. Thus, our approach allows us to enrich areas that fell within experimental ranges while directing the traditional CFU higher in order to fill out the whole experimental range.

### 2.4. Uncertainty analysis

LHS and partial rank correlation coefficients (LHS-PRCC) are used to perform an uncertainty analysis.(52) LHS-PRCC has been used in similar systems to characterize monotonic relationships between inputs and outputs.(52) One thousand samples are selected from the initial range using LHS and run with 7 replicates. These replicates are averaged before PRCCs are calculated at day 2 before the T cells are added and day 6. A significance level of 0.01 is used with a Bonferroni correction for the number of tests run. The relationship between the 50 varied parameters and 9 outputs of interest (totalMtbCount, mtbKilledByActivatedMacCount, mtbKilledByRestingMacCount, mtbKilledByCD8Count, activatedCD4Count, totalActivatedCD8s, activatedMacroCount, totalStat1MacroCount, totalNfkbMacroCount) are analyzed.

### 2.5. Matching Unpaired Runs

To be able to explore output spaces that are not accessible using the paired simulations described above, we also analyze matched simulations. Unpaired spheroid and traditional simulations are matched by selecting runs with similar (but not identical) initial condition parameters: *CD8Fraction*, *fractionCD4*, *fractionTBSpecific*, and *tbSpecificCD8Fraction*. To identify matched simulations, the spheroid runs are looped through for each traditional run, and a cost function was calculated. This function sums squared errors divided by maximum value for these 4 controlled parameters (*CD8Fraction*, *fractionCD4*, *fractionTBSpecific*, and *tbSpecificCD8Fraction*). The spheroid run with the lowest cost is selected to be matched to the unpaired traditional run.

## 3. Results

### 3.1. Results from multiple systems can be reproduced with one in silico framework

We first test whether or not the multiscale model can recreate the experimental data for bacterial fold change, cell count, and cell viability at day 6. Using the calibration method described above, parameter sets are identified whose output fit criteria for both spheroid and traditional data. (Figure 2a-c) These simulations give CFU fold change outputs that span most of the experimental range, except for the highest experimentally measured CFUs in the traditional cultures. Together this suggests we are able to recreate experimental data from multiple *in vitro* systems using the same sets of parameters (Appendix Figure 1) and the same model structure.

**Figure 2:**
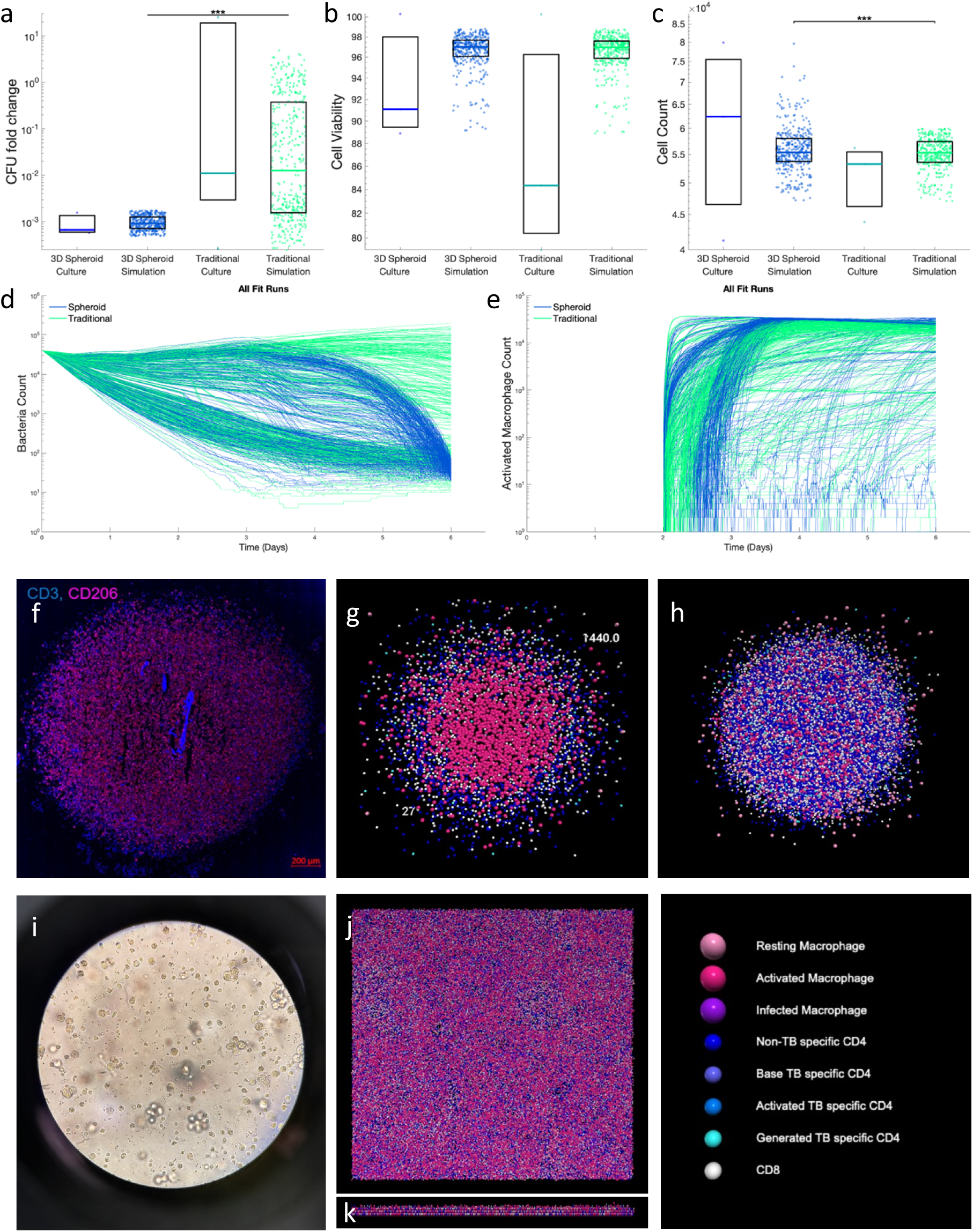
Paired simulations are calibrated to data from *in vitro* cultures. Spheroid and traditional simulations are run with the same parameters, only varying the initial spatial layout of cells and the movement rules. Comparison of experimental data to calibrated simulation data for a) CFU fold change from 4 hpi to 6 days, b) cell viability at day 6, and c) cell count at day 6. d) Bacterial count dynamics for calibrated spheroid and traditional simulations over the 6 day time course show heterogeneous behaviors. Spheroid and traditional simulations are visualized at day 6 for comparison to in vitro images. f) A slice of the *in vitro* spheroid culture on day 6. (Adapted with permission from Kotze et al. 2021) g) A slice through the center of a spheroid simulation. h) Full spheroid simulation. i) A brightfield image of the *in vitro* traditional culture on (day 6). j) Traditional simulation viewed top down. k) Traditional simulation viewed from side. *** p≤1e-3

After calibrating to both experimental systems, representative calibrated runs are visualized to compare with experimental images as a qualitative validation. Simulated spheroids (Fig. 2g,h) qualitatively match experimental microscopy (Fig. 2f), having a layered structure with macrophages on the inside and T cells in a cuff around the edge. The whole spheroid is situated in the middle of the simulated space with a very dense center with some cells less densely around the outside. The layered structure of the spheroid can be contrasted with the more well-mixed and dense traditional simulation (Fig. 2j,k) and experiments (Fig. 2i). These cells are localized at the bottom of the simulation space, due to the gravity-limiting spatial rules. These visualizations also highlight the versatility of the computational model, allowing the same base set of rules to recreate multiple *in vitro* culture systems. In summary, this quantitative calibration and qualitative validation indicates that our simulation-predicted spatial organization aligns well with experimental data.

Beyond recreating existing experimental data, our computational model can also predict high time resolution outcomes. Bacterial time courses show the heterogeneity of behaviors possible given both the initial conditions and the experimental range at day 6. (Fig 2d) This heterogeneity can give us insight into potential system dynamics and generate new testable hypotheses. Predictions can then be tested by designing experiments to distinguish among predicted behaviors by identifying time points and outputs of interest with the simulation. For example, macrophage activation (Fig 2e) could be compared with M1 activation markers *in vitro* at day 2.5 to differentiate between the two groups of spheroid simulations with different predicted timings of macrophage activation.

Taken together, these results indicate that that our computational framework can reproduce both bulk and spatial data from multiple experimental systems. Additionally, high time- and space-resolution predictions can be made about cell counts and interactions.

### 3.2. Our computational framework predicts high time resolution spatial data, including the evolution of a single spheroid over time

In *in vitro* and *in vivo* experiments, a granuloma must be destroyed to produce IHC or other data, meaning each time point corresponds to a different granuloma. In contrast, *in silico* models allow us to look at the evolution of spatial phenomenon *in situ*, meaning a single granuloma can be followed from creation to the end of the experiment.

These spatial dynamics can be analyzed both visually and quantitatively. Visually, an initial sphere of mixed infected and uninfected macrophages is seen at day 1 with a cuff of T cells being added at day 2 (Fig 3a). Macrophage activation starts at the interface of the macrophages and T cells and moves towards the center as time progresses. This activation corresponds to some T cell infiltration into the macrophage core. In this specific run, more CD8+ T cell activation leads to more infiltration by this population. Quantitatively, we can look at the radial density of cells and cell subpopulations to see similar trends (Fig 3b-e). Radial density graphs were generated by calculating the distances of the cells to the center of the spheroid, generating a histogram for the cells of interest by dividing them into preset bins, and then normalizing by the total volume in each bin which corresponds to the volume of a spherical shell. The simulation starts with uniformly distributed macrophages, before a cuff of uniformly distributed CD4+ and CD8+ T cells is added (Fig 3b). As time progresses the T cells spread out and begin to infiltrate the macrophage core, especially CD8+ T cells in this representative simulation. In our simulation, these T cells only contribute to the immune response when they are activated, so T cells are subdivided into resting and activated (Fig 3c). Activated T cells are more localized towards the center of the granuloma. This makes sense as interacting with a macrophage presenting antigenic peptides is required for T cell activation, and bacteria and macrophages that have interacted with bacteria are going to be localized to the core. These activated T cells provide one of the signals required for macrophage activation, STAT1 via IFN- γ. The distribution of this signal can be overlaid with the other required signal, NF-κB, giving insight into how macrophage activation propagates from the outside in (Fig 3d). The distribution of fully activated macrophages (Fig 3e) closely follow the STAT1 signal, suggesting in this model T cells are the limiting step of activation. The widespread NF-κB activation suggests it is not the limiting step, especially as macrophages are NF-κB activated from day 1 forward. This is likely due to TNF-α secretion from the intermixed infected macrophages.

**Figure 3:**
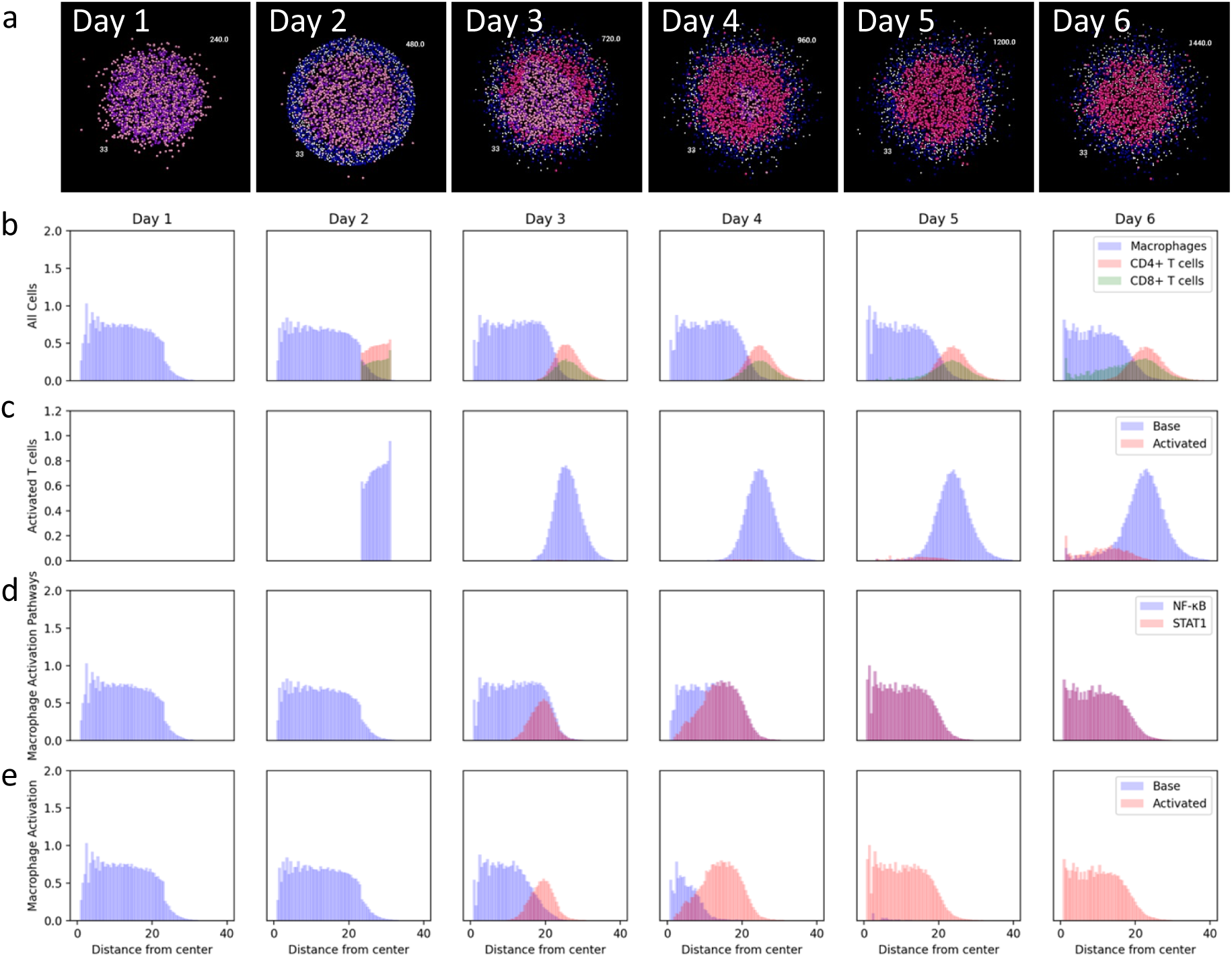
a) The spatial development of a single granuloma over 6 days. The radial distribution of b) macrophages, CD4+ T cells, and CD8+ T cells; c) base and activated T cells; d) NF-κB and STAT1 activated macrophages; e) base and activated macrophages.

Taken together, this illustrates how we can use our models to quantify key host- pathogen interactions in space and time in a single granuloma.

### 3.3. A distribution of outcomes, including T cell infiltration, is seen among spheroid simulations

Given that T cell signaling is important for macrophage activation, T cell infiltration becomes an output of interest. Visually, we noted the variation in T cell infiltration of the spheroids between parameter sets (Fig. 4a,b). Calibrated runs can show almost no infiltration (Fig. 4a) to almost homogeneous mixing of macrophages and T cells (Fig. 4b). To evaluate the heterogeneity across all of our simulations, the mean and standard deviation of radial density is calculated for macrophages and T cells for each simulation. This gives a mean position when correcting for the uneven volumes of the radial spheres. These measures for T cells and macrophages range from having nearly complete overlap to almost complete separation (Fig 4c). Over half of the simulations have macrophages with means around 10. However, many simulations also have higher macrophage means closer to the T cell means, suggesting more intermixing between T cells and macrophages among these runs. Little infiltration was seen in the *in vitro* model at day 6,(13) which aligns with some but not all of our simulated runs. Either the small sample size of the experimental study doesn’t account for full heterogeneity or this information can be used to further narrow the parameter space moving forward.

**Figure 4:**
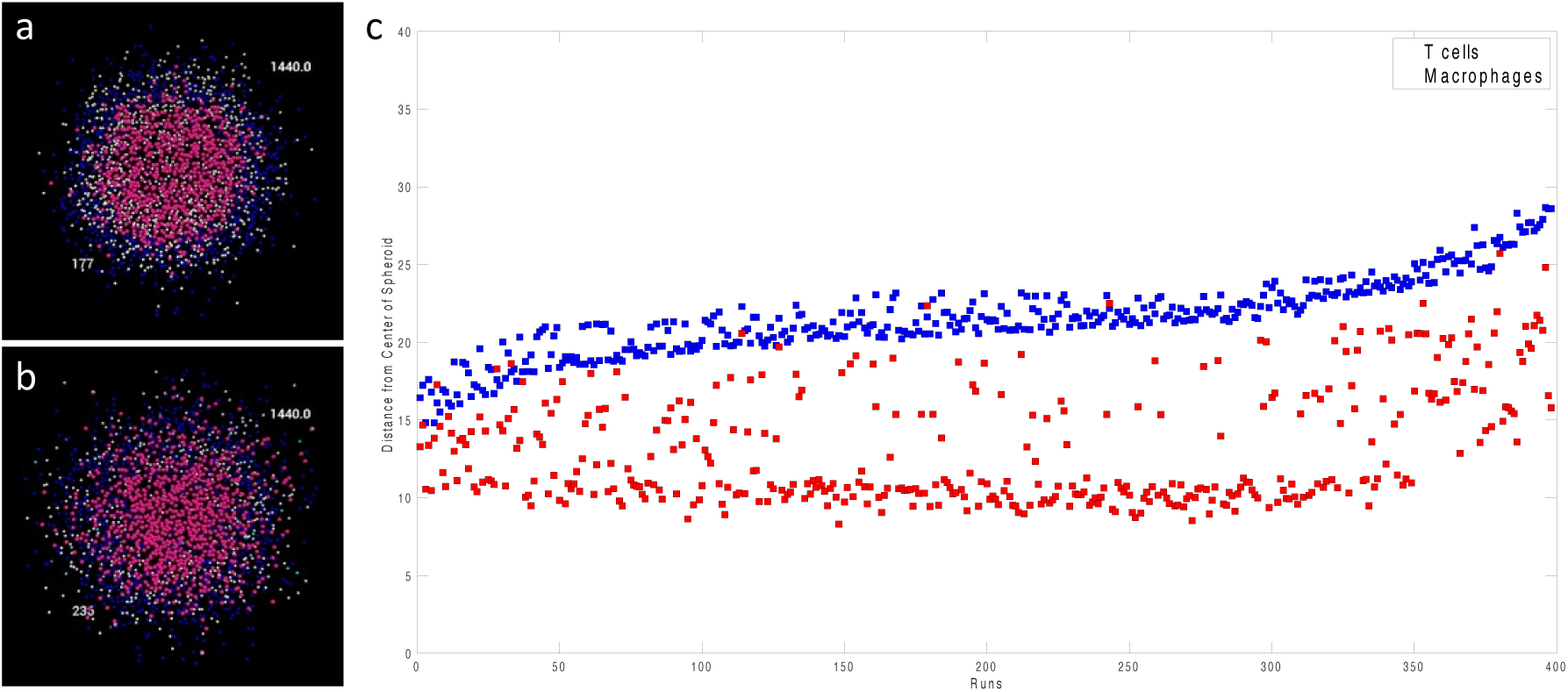
Example simulations with a) extensive and b) limited T cell infiltration into the macrophage core. A slice through the center of the granuloma is shown at day 6. c) Mean and standard deviation radial density of macrophages and T cells. Runs are sorted by mean T cell distance minus one standard deviation.

One way to look at the infiltration of T cells is to look at the difference between the T cell mean and macrophage mean. The higher the value the more separation between the cell types and, therefore, more structure. Spearman’s rank correlation coefficients were calculated between this distance measure of separation and outputs of interest at day 6 with α = 0.01. Our model suggests this measure of separation is not significantly correlated to total bacterial count (ρ = 0.091292, p = 0.068859). However, this model is only looking at runs that were calibrated to experimental data, which has a small range of bacterial count for the spheroid simulations. If increasing T cell separation was an isolated change it’s possible that this relationship would be seen.

Although the bacterial counts in the spheroid simulations are all within a small range, the bacterial counts in the corresponding traditional simulations vary more. The Spearman’s rank correlation coefficient between our separation measure and the traditional total bacterial count shows a significant positive correlation (ρ = 0.561759, p < 0.000001). The parameter sets that show more separation in the spheroid have higher bacterial load in the traditional cultures. So, this would suggest that those parameter sets rely on a lot of structure to be able to control bacteria, because when those parameters are used to simulate the traditional well-mixed conditions, the bacteria are not as well controlled. On the other hand, those parameter sets that don’t have a lot of separation, do equally well in controlling bacteria in both the spheroid and traditional. Parameter sets with less separation resemble the traditional organization, so similar results are expected. So, bimodal results are seen, where there’s two different ways that the model can control the bacteria – one is structure dependent and the other is not.

Other correlations seen between separation measure and outputs of the spheroid model include *activatedMacroCount* (ρ = 0.307164, p < 0.000001), *activatedInfectedMacroCount* (ρ = 0.419979, p < 0.000001), *totalStat1MacroCount* (ρ = 0.307557, p < 0.000001), and *totalActivatedCD8s* (ρ =0.171726, p = 0.000580). Increasing separation correlates to more STAT1 and total macrophage activation, including activation of the infected macrophage population. CD8+ T cell activation is also positively correlated with separation. Due to the distribution of separation, we are able to look at the impact of spatial layout of cells on many outputs.

Taken together, these results illustrate that our simulations can produce a wide range of outcomes that are consistent with the experimental data; and how these simulations can be used to explore how granuloma structure impacts bacterial control and activation.

### 3.4. Comparisons between the models using uncertainty analysis can help identify ideal use cases

LHS-PRCC is performed on the initial large LHS sweep to quantify how uncertainty in the parameters impacts uncertainty in the outputs of both the spheroid and traditional simulations. At day 2 before the T cells have been added to the simulation, the spheroid and traditional simulations show similar responses to changes in parameters. Total bacterial count is inversely correlated with the doubling time of the internalized bacteria and the killing ability of the resting macrophages. All of the bacteria at the beginning of the simulation are internal due to a washing step after the infection of the macrophages. Lower doubling times of these bacteria lead to more generations and more bacteria. Poorer killing ability of the resting macrophages leads to more bacteria.

Before the addition of T cells, resting macrophages are responsible for all of the bacterial killing. Cytotoxic CD8+ T cells have yet to be added to the culture, and T cells are required to fully activate macrophages. So, *mtbKilledByRestingMacCount* accounts for all of the killing and closely aligns with the *totalBacterialCount*. Base killing probability is positively correlated with *mtbKillingByRestingMacCount*, as better killing ability leads to more killed bacteria.

Total NF-κB activated macrophages is the only other output showing significant correlations with parameters before day 2. NF-κΒ signal can come from TNF-α or bacteria and is required for total activation. Total NF-κB activated macrophages are correlated to *nfkbSpan*, *baseKillingProbability*, *InfectedMacrophageTNFSecretion*, and *macrophagePopulation_MaxLifespan. nfkbSpan* is the length of time that NF-κB stays active after receiving the initial signal, so the longer this time period is the more NF-κB activated macrophages there are. When *baseKillingProbability* is lower, fewer bacteria are killed, and more bacteria and infected macrophages are available to activate NF-κB. Higher TNF-α secretion from infected macrophages also leads to more activation. Lastly, longer macrophage lifespans mean more macrophages are alive to be activated.

After the T cells are added, the total bacteria count is more dependent on CD4+ T cell parameters. The internal doubling time and base killing probability are both still negatively correlated with total bacteria count. The rest of the significantly correlated parameters are associated with CD4+ T cells and STAT1 activation. Fewer TB specific CD4+ T cells, less CD4+ T cell activation, and more CD4+ deactivation all reduce the amount of activated CD4+ T cells indirectly leading to more bacteria. Higher threshold for STAT1 activation by IFN-γ, higher degradation rate for IFN-γ, and less CD4+ T cell IFN-γ secretion all lead to less macrophage STAT1 activation. Again, this will indirectly lead to more bacteria. With the inclusion of adaptive immune cells, the responses of the two set ups also diverge more. For bacterial counts the only difference is due to external bacteria. Lower external doubling time leads to more bacteria only in the traditional simulation, as the population of external bacteria is so small in the spheroid simulation.

Macrophage activation is mostly dependent on CD4+ T cell parameters in both models. However, increased macrophage activation in the traditional simulation is also correlated with increased macrophage lifespans and decreased CD4+ T cell doubling time. This suggests that macrophage death might be limiting the population size in traditional runs. Also, a higher CD4+ T cell population will lead to more macrophage activation.

CD8+ T cell activation is correlated with parameters related to CD4+ T cells, CD8+ T cells, and IFN-γ. Differences between the two models include negative correlations between the probability of deactivation/population doubling time and total CD8+ T cell activation. Less deactivation or more proliferation should lead to higher activated populations.

The only parameter-output relationship seen just in the spheroid simulation is a positive correlation between the *baseKillingProbability* and total NF-kB activated macrophages. This is the opposite of the relationship seen at day 2. One hypothesis for this relationship is that more killing initially leads to less macrophage activation and subsequent death. This is supported by positive correlations between macrophage lifespans and this output.

Despite these differences, the two models have many similar responses to changing parameters. For example, the total activated CD4+ T cells show the same relationships for both the spheroid and traditional simulations with regards to all parameters. The significantly correlated parameters are all related to CD4+ T cells: fraction of TB specific cells, activation probability, deactivation probability, and doubling time.

Altogether, these results show that similar parameters are driving dynamics in the spheroid and traditional models before day 2, but the influential parameters diverge after the addition of T cells. These correlations can be used to select what *in vitro* model is needed when designing experiments, as main drivers of outputs can be identified. For example, the traditional simulation has a correlation between macrophage lifespans and macrophage activation that is not seen in the spheroid. This relationship suggests that macrophage lifespans influence macrophage activation in the traditional culture, so a spheroid might be more appropriate if the biological question under investigation relates to drivers of macrophage activation.

### 3.5. Limitations in representing both systems can guide future model iterations

The analysis done thus far is based on paired calibration. Pairing the simulations makes the assumption that everything except for the initialization and movement is the same between the spheroid and traditional simulations. While this assumption allows us to recreate a majority of the experimental range, the highest traditional CFU counts are unable to be recreated with paired runs. Traditional runs with high levels of bacteria falling in this range are seen, but the corresponding spheroid simulation did not meet calibration criteria. This suggests that in order to reproduce these high traditional CFU results, some parameters (i.e. biological mechanisms) may need to be different between the spheroid and traditional simulations. To investigate this possibility, we evaluate unpaired simulations that are allowed to have different parameter values between traditional and spheroid simulations, but that are matched as closely as possible for initial conditions that are expected to be the same.

Traditional runs are matched with spheroid runs that meet calibration criteria and have similar initial conditions as defined in the methods. After these runs are matched, the parameters of the spheroid and traditional simulations are compared. Nine parameters are found to be significantly different (Figure 6). Some of these parameter differences can lead to less bacteria in the spheroid directly or indirectly by increasing activation. The matched spheroid runs had higher internal doubling time of the bacteria meaning the bacteria grow more slowly and a higher resting macrophage killing rate leading to more bacterial killing. Therefore, the matched spheroid runs have less bacteria than the traditional runs directly due to less growth and more killing. The matched spheroid runs also have parameter differences that lead to more macrophage activation. Lower *IFNthresholdForStat1Activation* in spheroid runs would give more STAT1 activation of macrophages causing more overall macrophage activation. Lower *CD4DeactivationProbability* in spheroid runs would prolong CD4+ T cell activation giving these cells more opportunities to activate macrophages. Lastly, lower *TNFDegradationRatePerSecond* in spheroid runs maintains higher concentrations of TNF-α, leading to more NF-κB activation of macrophages.

**Figure 5:**
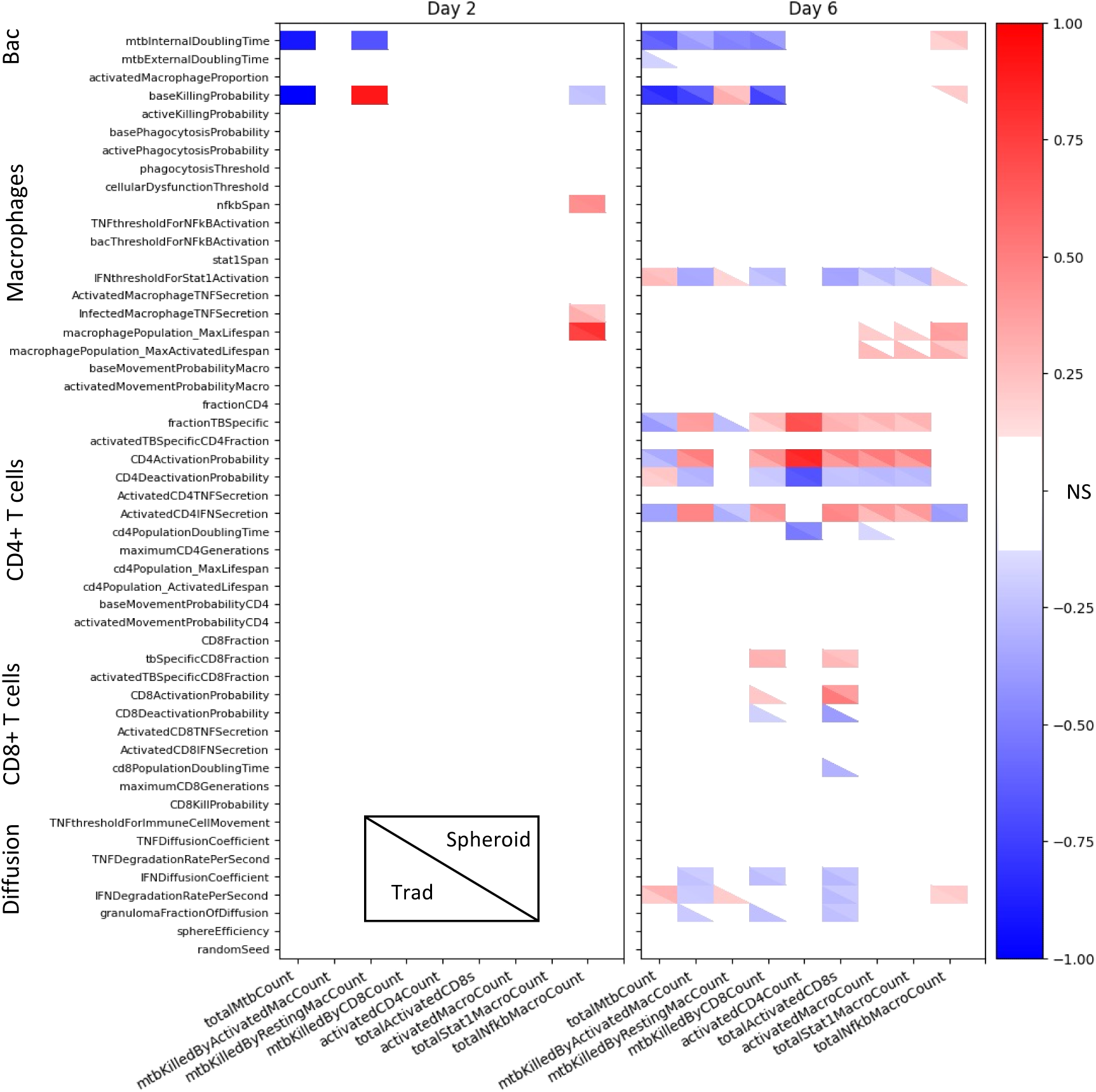
Impact of input parameters on simulation outputs at day 2 before the T cells are added and day 6. Correlation coefficients for spheroid and traditional simulations are shown in the same heatmap with the traditional in the lower left hand corner and the spheroid in the upper right. Insignificant correlations are shown in white, while positive and negative correlation are shown with red and blue, respectively. Significance was determined with α =0.01 and a Bonferroni correction.

**Figure 6:**
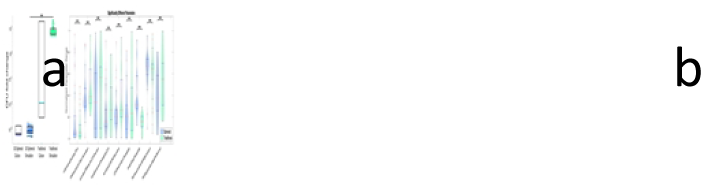
a) Traditional simulations that fell above the paired range and matched spheroid simulations. b) Significantly different parameters between the set of high traditional simulations and matched spheroid simulations.

The role of the other parameters is less clear. Lower *activatedTBSpecificCD4Fraction*, lower *ActivatedCD8TNFSecretion*, and higher *TNFthresholdForNFkBActivation* would all suggest lower macrophage activation in the spheroid. Lower *baseMovementProbabilityCD4* could delay activation of TB specific T cells or could lead to less spatial interference by non-TB specific T cells in spheroids. As these simulations are matched after the fact, some of these differences are potentially spurious. However, these differences can guide future computational and experimental studies by highlighting hypothesized functional differences between traditional and spheroid cultures.

## 4. Discussion

*In silico* models have been used previously to represent multiple *in vitro* systems for other diseases. In 2006, Grant et al. used cellular automata to represent the growth of epithelial cells in 4 conditions: 3D embedded, suspension, surface, and collagen overlay cultures.(62) They were able to recreate the complex structure associated with each condition with a set of axioms governing the interactions of cells, matrix, and cell-free space. The difference between a 2D and 3D culture system has also been modeled to explore viral dynamics and drug toxicity. A network model of tumor cell infection by oncolytic viruses was simulated in a 2D monolayer and 3D environment.(63) This model suggested that traditional mean field models overestimate how effective therapy would be. Beyond this, infection in a 3D environment was shown to have a smaller chance of tumor eradication, emphasizing the need for ideal virus characteristics: fast replication and slow tumor cell killing. A virtual cell based assay was extended from 2D cultures to 3D spheroids to predict drug toxicity.(64) This model was found to represent 3D *in vitro* models well, which show higher drug toxicity than 2D monolayers.

In all instances, space is explicitly modeled to gain insight into the system behavior in different configurations. The spatial configurations alter dynamics of the system and can change important predictive outcomes, such as drug response. Similar to these prior works, we explicitly include space to model two different environmental setups. We show that spatial organization alone can change the dynamics of the system and primary outcome, bacterial count. Moving forward, we can use our simulations to predict which model outcomes are likely to be affected by spatial organization and therefore guide experimental decisions.

Note, we are using a single *in silico* framework to represent separate traditional and 3D cell cultures. A separate problem, representing one *in vivo* or *in vitro* system with both a 2D and 3D computational model, has also been addressed.(65, 66) Models of *in vivo* granulomas and *in vitro* spheroids suggest that 2D representations of 3D systems (i.e. slice through center of structure) have similar results and save computational time.(65, 66)

Granulomas are spatial organized structures, with a core of macrophages and a cuff including CD4+ and CD8+ T cells.(67) The center of the granuloma is a more pro-inflammatory environment, while the cuff has more anti-inflammatory cytokines.(68, 69) Higher frequencies of pro-inflammatory cytokines or lymphocytes are correlated with lower bacterial burden, but it is suggested that a balance of pro- and anti-inflammation is necessary to limit both bacterial growth and pathology.(69–71) While we don’t explicitly include anti-inflammatory pathways in this preliminary model, our model does show pro-inflammatory signals localized to the core. Specifically, macrophage activation is limited by the interactions between IFN-γ and macrophages which begins at the periphery of the core and moves inwards. This looks similar to the pattern of p-STAT1 seen in peripheral regions in immunohistochemistry of NHP granulomas.(72) Previous computational modeling also suggests the importance of IFN-γ producing T cells and interactions between macrophages and T cells for bacterial control.(73) It emphasizes the importance of spatial organization as interactions between CD11c+ macrophages and T cells are limited due to the cellular distributions within granulomas and the recruitment of non-specific T cells.

While the structure of our model is artificially constructed rather than emerging from immune interactions, the similar spatial patterns for cells and activation is encouraging. Further comparison could be accomplished by applying our methods for analyzing the distribution of cell types and signals within granulomas to *in vitro* and *in vivo* data in the future.

The evolution of a single granuloma can be followed over time in other systems. Sequential imaging with [18F] fluorodeoxyglucose positron emission tomography and computed tomography has been used to follow disease progression in NHP and track response to TB treatment in humans.(74–76) This imaging gives information at the lesion-tissue scale. Florescent *in vivo* microscopy of zebrafish embryos has given insight into the cellular level dynamics.(77) Imaging after infection of zebrafish embryos with *Mycobacterium marinum* allows tracking of infected macrophages providing information about early granuloma formation and dissemination. Recently, a method to study zebrafish granulomas *ex vivo* called Myco-GEM was created that allows continuous lightsheet imaging for upwards of 8 hours.(78) With tagging of cytokines, specific cells, or bacteria the inflammatory state of the granuloma, granuloma dynamics, cell movement, and bacterial load can be longitudinally examined.

Our model similarly provides dynamic information at the cellular scale. Beyond this, we can gather information about bulk cell counts, cell activation status, and cytokine concentrations without perturbing the observed system. Thus, our computational model can complement *in vitro* experimental systems, by providing both high-resolution spatiotemporal information and bulk information about host-pathogen interactions within individual granuloma structures. Simulations with virtual perturbations on knockouts can then quickly be run to examine how these interactions contribute to bacterial survival or elimination.

TB is a very heterogenous disease. There are many different clinical outcomes: bacterial clearance, asymptomatic latent infection, and active infection.(71, 79) These host level outcomes are dependent on a population of granulomas, which can be very heterogenous even within the same lung.(76, 80)Granulomas have many different structures which can lead to bacterial dissemination, control, or clearance.(71, 79) Our *in vitro* models are more controlled with an established structure and proportion of cell types. Smaller sample numbers still showed a large range of bacterial control, which can be recreated *in silico*. We also see heterogeneity in T cell localization *in silico*. While this is not seen as much *in vitro*, there is some variability *in vivo*. Early granulomas have T cells dispersed throughout, while well-developed ones are more structured with a ring of T cells.(81) Being able to reproduce a diversity of granuloma organizations will allow us to explore how different microenvironments contribute to granuloma trajectory and treatment response.

LHS-PRCC has been used to look at correlations between inputs and outputs in simulations of *in vivo* NHP granulomas. While our time points don’t line up with the longer *in vivo* simulations, we can compare parameter influences before and after adaptive immunity has been added. In the first iteration of the NHP granuloma simulation, there are similarities to our model.(19) This model from literature shows a strong positive correlation between intracellular growth rate and total extracellular bacteria during early infection.(19) As infection progresses extracellular bacteria in the simulated NHP granulomas becomes negatively correlated with T cell parameters, namely recruitment, movement, and activation of macrophages.(19)

In these simulations of *in vivo* granulomas all bacteria start extracellularly, while all bacteria start intracellularly in our *in vitro* model. Our primary output of interest then becomes intracellular bacteria, which shows a similar relationship with the intracellular growth rate before the addition of the adaptive immune system. Some comparisons between this *in vivo* simulation and our *in vitro* simulation are limited because *in vivo* mechanisms are missing *in vitro,* like cellular recruitment. But we see an increased importance of T cell parameters on our output of interest after adaptive immunity is initiated as in this literature model. In our short term ‘artificially’ assembled spheroids, we see similar parameter influences to *in vivo* granulomas. Therefore, some comparisons can be made not only between our two *in vitro* models, but also *in vivo* simulations, to be able to rationally identify good use cases for various *in vitro* systems.

Our model is not without limitations. The only PBMC derived CD3+ T cells simulated are CD4+ and CD8+ T cells. Some subsets of T cells (e.g. regulatory and γδ) are excluded from the model for the purpose of simplification. Simplifications are also made to the macrophage activation pathway. The model only incorporates M1 macrophage polarization/activation represented as a 2-step pathway, and M2 macrophage polarization is not included. Additionally, our model has been calibrated to be used with cells from patients with presumed active TB. The exact same cells derived from an uninfected patient or a patient with latent TB might behave differently, and the model would need to be recalibrated to different data. These assumptions can be reassessed as we iterate this model to use it in answering new biological questions.

While our model is able to represent a majority of the characteristics that could be incorporated into a complex *in vitro Mtb* model, it still diverges from the idealized model in a couple ways. No explicit environment impact on the cells in the simulation is included. It’s known that plastic and glass plates differ from *in vivo* environments, and as such extracellular matrix (ECM) components like collagen have been incorporated into *in vitro* models. ECM can also change the lifespan and movement of the cells and sequester chemokines. We plan to incorporate ECM in future iterations. While primary human cells were represented, the bacteria represented within this model is BCG, a model organism for *Mtb*, rather than *Mtb* itself. BCG was used for preliminary analysis as it can be used outside of a BSL3 laboratory. Switching between BCG and *Mtb* could be done by adjusting parameter values, but more detailed pathways would need to be added if specific virulent strains were of interest.

## 5. Conclusion

In summary, we show a novel application of ABMs to *in vitro* TB infection culture systems. In doing so, we introduce a framework to potentially integrate results from and compare multiple *in vitro* models.

## Acknowledgements

This work used the Extreme Science and Engineering Discovery Environment (XSEDE), which is supported by National Science Foundation grant number ACI-1548562. Anvil at Purdue and Expanse at UCSD were used through allocation TG-MDE220002. We also thank Lev Gorenstein and the rest of the Research Computing Staff for their assistance with batch computing at the Rosen Center for Advanced Computing. We would also like to acknowledge Catherine Weathered for her mentorship and her work setting up the foundations in Repast and Slurm for our lab.

## Supplementary Information Captions

**Table 1:** Parameters that were held constant during sampling, their values, and units.

**Figure 1:** Distribution of parameters in calibrated runs. The ranges of the parameters have been normalized from 0 to 1 with the bounds representing the minimum and maximum of the ranges listed in *Table 1*.

